# Pre-Omicron Immunity Generates IgG⁺ but Not IgA⁺ Memory B Cells Reactive to Omicron Spike Protein

**DOI:** 10.1101/2025.09.26.678818

**Authors:** T Tsoleridis, I Singh, D Onion, JG Chappell, A Kelly, J Nightingale, AM Valdes, BJ Ollivere, KB Hoehn, RA Urbanowicz, JK Ball

## Abstract

The emergence of the Omicron variant marked a major antigenic shift from previous SARS-CoV-2 variants. While the emergence of new variants of concern was associated with loss of protection from infection, protection from serious disease was maintained. What was less clear was whether this protection was associated with expansion, affinity maturation of pre-existing cross-reactive B cells, or *de novo* generation of variant-specific B cells. To define the cross-reactivity of memory B cell populations generated through infection and/or vaccination with pre-Omicron variant spike, we utilised archived samples from healthcare workers enrolled in the extensively monitored Panther study cohort. Using samples collected from Omicron-naïve individuals approximately three months (106 days) after a third BNT162b2 vaccine dose, we analyzed the spike-specific B cell repertoire via multiparameter flow cytometry and B cell receptor (BCR) sequencing.

Reactivity to both Lineage A and Omicron spike proteins was higher for serum IgG than IgA, and serum neutralisation potency was significantly reduced against pseudoviruses bearing the Omicron spike compared to those with Lineage A spike. The frequency of Omicron-reactive IgA+ B cells was markedly lower than that of Lineage A-reactive cells, whereas Omicron-reactive IgG+ memory B cells were more frequent. These responses were predominantly localized to classical memory and double-negative (DN) B cell subsets. BCR sequencing of donors unexposed to Omicron confirmed that Omicron-binding clones were class-switched and somatically mutated, indicating they originated from pre-existing memory B cells primed by vaccination or infection.

Our findings demonstrate that memory B cells generated through infection and/or immunisation with pre-Omicron variants were primed for broad antigenic recognition despite Omicron’s extensive genetic divergence. However, the diminished IgA+ B cell frequency suggests a weakened mucosal antibody defence, potentially explaining the continued susceptibility to infection despite preserved protection from severe disease.

## Introduction

Since the emergence of SARS-CoV-2, there have been several variants of concern (VOC) that spread rapidly, displacing prior variants. In November 2021, Omicron was identified as a new VOC [1]. Omicron has 37 amino acids substitutions in the S protein, 15 of which are in the receptor binding domain (RBD), compared to Lineage A. Studies have shown that it replicates faster in bronchi but less efficiently in the lung parenchyma and is more dependent on cathepsins rather than TMPRSS2, suggesting that it favours replication in the upper respiratory tract compared with the other variants [2]. The higher infection rate could be attributed to the ability of Omicron to escape neutralising antibody responses, with individuals showing a ∼20-fold reduction in vaccine-elicited antibody neutralisation against this variant [3–7]. Despite this reported reduction in cross-protection by serum neutralising antibodies, research has shown that two or three-dose vaccination still protected individuals infected with Omicron against severe disease and hospitalisation, although protection against infection and transmission was reduced [8–10]. Importantly, studies have suggested that previous infection or vaccination with spike from earlier SARS-CoV-2 variants can induce cross-reactive T-cell and B-cell memory responses, which protect from severe COVID-19 [11–16].

Studies showing Omicron escape in individuals vaccinated with Lineage A are based on serum circulating antibodies which may not fully reflect the memory compartment. To better understand the relationship between serum antibody and memory B cell populations, and their potential importance in protection from infection or disease, we used serological and multi-parameter flow cytometry assays. This allowed us to delineate the Lineage A and Omicron-specific antibody response in a cohort of vaccinated healthcare workers, some of whom had documented pre-Omicron SARS-CoV-2 infection [17]. Our data shows that the Omicron-reactive peripheral IgG^+^ B-cell populations are present in vaccinated individuals, both previously infected and uninfected, prior to any exposure to the Omicron variant.

## Results

### Serological responses to Omicron are present, but reduced, following vaccination with or without non-Omicron infection

Eighteen (n=18) vaccinated healthcare workers who were part of the Panther cohort study [17] were recruited, twelve (n=12) of whom had had a previous SARS-CoV-2 infection and six (n=6) had not (**Figure 1A**). Sixteen donors had previously received two initial doses and one booster of the BNT162b2 (Pfizer) vaccine and two had received two initial doses of ChAdOx1-S (AstraZeneca) and a booster of Pfizer. The first dose was administered in December 2020/January 2021, the second in February/March 2021 and the booster in September/October 2021. EDTA blood samples were collected in January 2022, approximately 106 days (ranging from 70 to 126 days) since the donors received their booster dose, to assess their B cell and antibody responses against Lineage A and Omicron variants (**Table 1**). Critically, all donors were vaccinated before the emergence of the Omicron variant and had no evidence of infection with Omicron between vaccination and sample collection. Thus, any Omicron reactivity observed was induced from either vaccination with Lineage A and/or infection with a variant other than Omicron.

**Figure 1.**
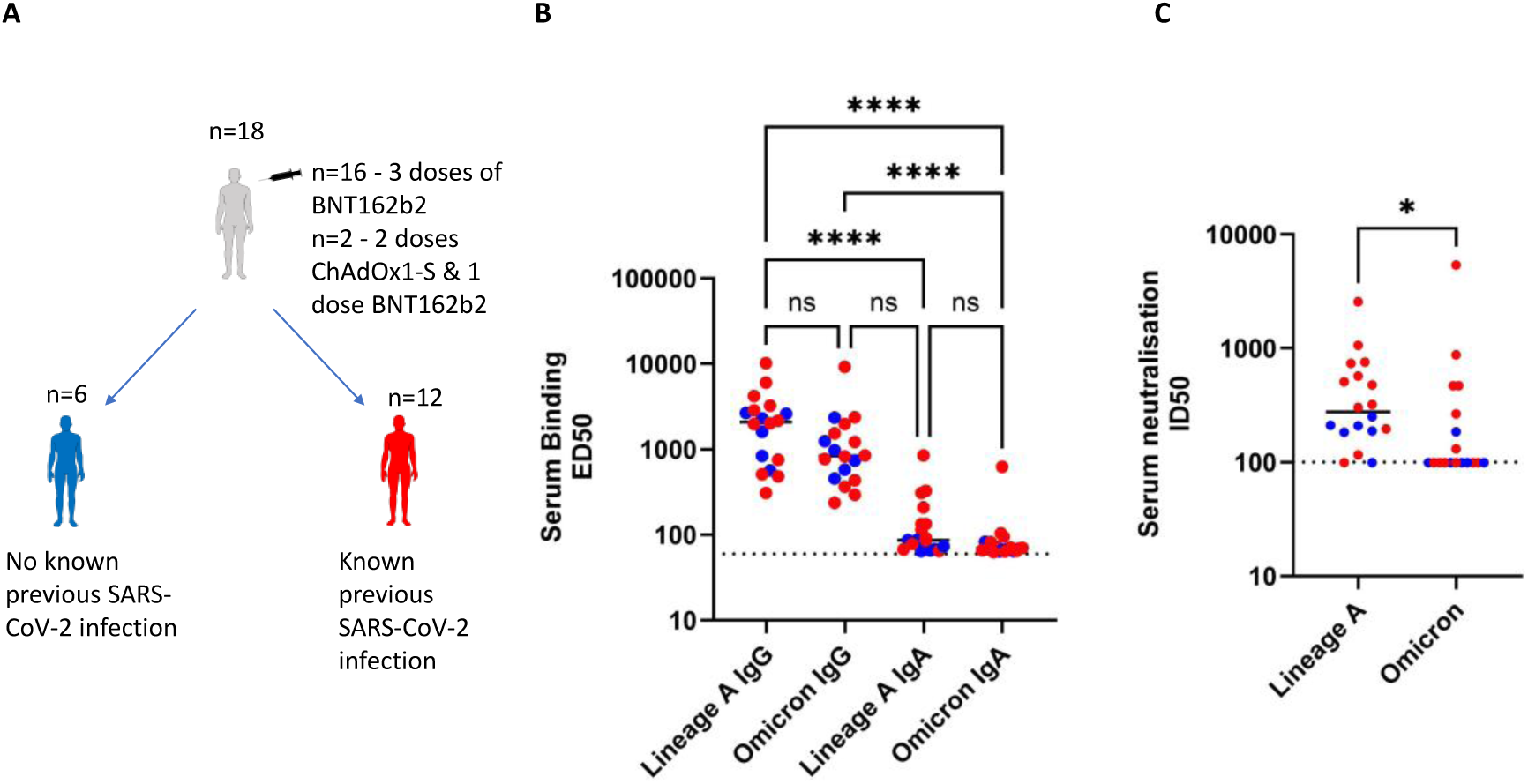
Reactivity and neutralising ability of serum antibodies (**A**) Graphical summary of the (n=18) healthcare workers who participated in the study; n=16 of them have had 3 doses of BNT162b2 and n=2 have had 2 doses of ChAdOx1-S & 1 dose BNT162b2, (blue, n=6) have not had recent known SARS-CoV-2 infection whereas (red, n=13) had. (**B**) ED50 plot of uninfected (n=6, blue) and previously infected (n=12, red) donor serum IgG and IgA binding against Lineage A and Omicron variants showed no significant differences in reactivity between Lineage A and Omicron for neither IgG nor IgA. Significant differences were observed between Lineage A IgG – Lineage A IgA (p<0.0001), Lineage A IgG – Omicron IgA (p<0.0001) and Omicron IgG – Omicron IgA (p<0.0001) (**C**) ID50 plot of uninfected (n=6, blue) and previously infected (n=12, red) donor serum against Lineage A and Omicron pseudotypes showed significantly higher (p = 0.025) neutralisation potency against Lineage A compared to Omicron. ID50 values < 100 are presented as ID50 value 99. All statistical tests were performed using Prism 9.0 [(**B**) Friedman paired Anova & Dunn’s multiple comparison tests (**C**) Wilcoxon paired two-tailed t-test].

**Table 1.** Demographics of donors. First 12 have a history of infection and the last six do not. Infection status obtained through Roche N ELISA. All donors were also testing weekly with a lateral flow device.

| Donor | Gender | Manufacturer primary dose | 2nd dose Days after initial | Booster Days after initial | Infection status Jul 2020 | Infection status Sept 2020 | Infection status Mar 2021 | Infection status Jul 2021 | Infection status Jan 2022 | Sample Days after booster |
| --- | --- | --- | --- | --- | --- | --- | --- | --- | --- | --- |
| 003 | F | Pfizer | 70 | 291 | Negative | Negative | Positive | Positive | Positive | 96 |
| 009 | M | AstraZeneca | 79 | 281 | Positive | Positive | Positive | Not done | Positive | 95 |
| 012 | M | Pfizer | 67 | 324 | Positive | Positive | Positive | Positive | Positive | 70 |
| 025 | M | Pfizer | 70 | 303 | Positive | Positive | Positive | Not done | Positive | 101 |
| 040 | F | Pfizer | 70 | 292 | Positive | Positive | Positive | Positive | Positive | 108 |
| 116 | M | Pfizer | 70 | 271 | Positive | Positive | Negative | Negative | Negative | 108 |
| 199 | M | Pfizer | 73 | 295 | Positive | Positive | Positive | Not done | Positive | 108 |
| 272 | F | Pfizer | 70 | 269 | Negative | Negative | Positive | Positive | Positive | 109 |
| 286 | F | Pfizer | 70 | 260 | Positive | Positive | Positive | Positive | Negative | 116 |
| 350 | F | Pfizer | 70 | 289 | Positive | Positive | Negative | Negative | Negative | 115 |
| 418 | M | Pfizer | 75 | 285 | Positive | Negative | Negative | Not done | Negative | 126 |
| 434 | F | Pfizer | 97 | 299 | Not done | Not done | Positive | Positive | Positive | 121 |
| 022 | M | AstraZeneca | 88 | 500 | Negative | Not done | Negative | Not done | Negative | 81 |
| 049 | F | Pfizer | 70 | 288 | Negative | Negative | Negative | Negative | Negative | 115 |
| 194 | M | Pfizer | 63 | 278 | Negative | Negative | Negative | Negative | Negative | 82 |
| 298 | M | Pfizer | 70 | 296 | Negative | Negative | Negative | Negative | Negative | 110 |
| 311 | F | Pfizer | 71 | 275 | Negative | Negative | Negative | Negative | Negative | 118 |
| 378 | F | Pfizer | 70 | 286 | Negative | Negative | Negative | Not done | Negative | 122 |

Enzyme-linked immunosorbent assay (ELISA) was performed to measure the peripheral IgG and IgA reactivity against Lineage A and Omicron trimeric spike. The reciprocal half maximal effective concentration (ED_50_) values for IgG against Lineage A and Omicron trimeric spike in had a median of 2,084 (308.7 – 10,179) and 837.7 (236.1 – 9,259) respectively. The ED_50_ values for IgA against the same spikes had a median of 86.54 (64.31 – 846.7) for Lineage A and 69.81 (62.58 – 625.4) for Omicron. Approximately 10-fold higher overall reactivity was observed in IgG compared to IgA (**Figure 1B**). Comparison of ED_50_ values did not show any significant differences in reactivity between the variants in either antibody isotype or sub-cohort.

Having shown that the overall serum antibody reactivity to Lineage A and Omicron trimeric spike were not significantly different, we next determined if there were differences in their ability to neutralise lentivirus pseudotypes supplemented with either Lineage A or Omicron spike. The median reciprocal half-maximal inhibitory concentration (ID_50_) value for Lineage A and Omicron pseudotypes were 276.5 (99 – 2,556) and 99 (99 – 5,388) respectively. Comparison of ID_50_ values showed a significantly lower (p = 0.025) neutralisation potency against Omicron pseudotypes (**Figure 1C**). These findings suggest that although serum antibodies bind equally well to Lineage A and Omicron trimer spike, they cannot neutralise Omicron pseudotypes with the same potency as Lineage A. (**Figure 1C**).

### Following vaccination against Lineage A, the frequency of spike-reactive peripheral IgA^+^, but not IgG^+^, B-cells was significantly lower against Omicron compared to Lineage A

Analysis of serum antibody responses showed a general trend towards lower neutralisation potency to the Omicron variant. However, serum antibodies are mainly produced by long-lived antibody secreting cells resident in the bone marrow. To see whether this general decrease in Omicron neutralisation was mirrored in Omicron-reactive circulatory memory B cell frequency we collected peripheral blood mononuclear cells from each donor and performed multi-parameter flow cytometric analysis using labelled trimeric spike of each variant. Surface markers were used to discriminate IgG^+^, IgM^+^, IgD/M^+^ and IgA^+^ expressing B cells and determine the frequency of B cells reactive to Lineage A and Omicron trimeric spike. The median frequency of IgG^+^, IgM^+^, IgD/M^+^ and IgA^+^ B-cells reactive to Lineage A and Omicron spike from total live B cells was 0.0997% & 0.0791%, 0.0015% & 0.0015%, 0.0118% & 0.0051% and 0.0518% & 0.0032% respectively. The frequency of Omicron reactive IgA^+^ B cells was lower than Lineage A spike-reactive B cells, and this difference was statistically significant (p = <0.0001) (**Figure 2A**). There were no significant changes in the frequencies of the other antibody isotypes. Moreover, three previously infected and one uninfected donor showed no IgM^+^ reactivity against Lineage A spike. Also, two infected and three uninfected donors showed no detectable IgM^+^ reactivity against Omicron spike. One donor from each group showed no detectable IgM^+^ reactivity against either spike (**Figure 2A**). To measure and compare the relative proportion of spike reactive B-cells for each antibody isotype, we calculated the Omicron:Lineage A ratio. The median Omicron:Lineage A ratio of IgG^+^, IgA^+^, IgM^+^ and IgD/M^+^ was 0.8296, 0.0425, 0.3892 and 0.4814 respectively. Comparison of the Omicron:Lineage A spike reactive B-cell ratio showed that IgA^+^ sub-populations were much more likely to bind to Lineage A than Omicron than IgG populations (p<0.0001) (**Figure 2B**). This shows that the proportion of Omicron-reactive IgA^+^ compared to IgG^+^ B-cells was significantly lower. Similarly, the ratios for IgM^+^ and IgD/M^+^ were significantly higher than IgA^+^(p = 0.0029 & p = 0.0059 respectively). Therefore, despite broadly comparable memory B cell responses to SARS-CoV-2 Lineage A and Omicron across most isotypes, Omicron-specific IgA+ B cells were significantly reduced in frequency, suggesting a possible deficit in mucosal B cell immunity against the Omicron variant.

**Figure 2.**
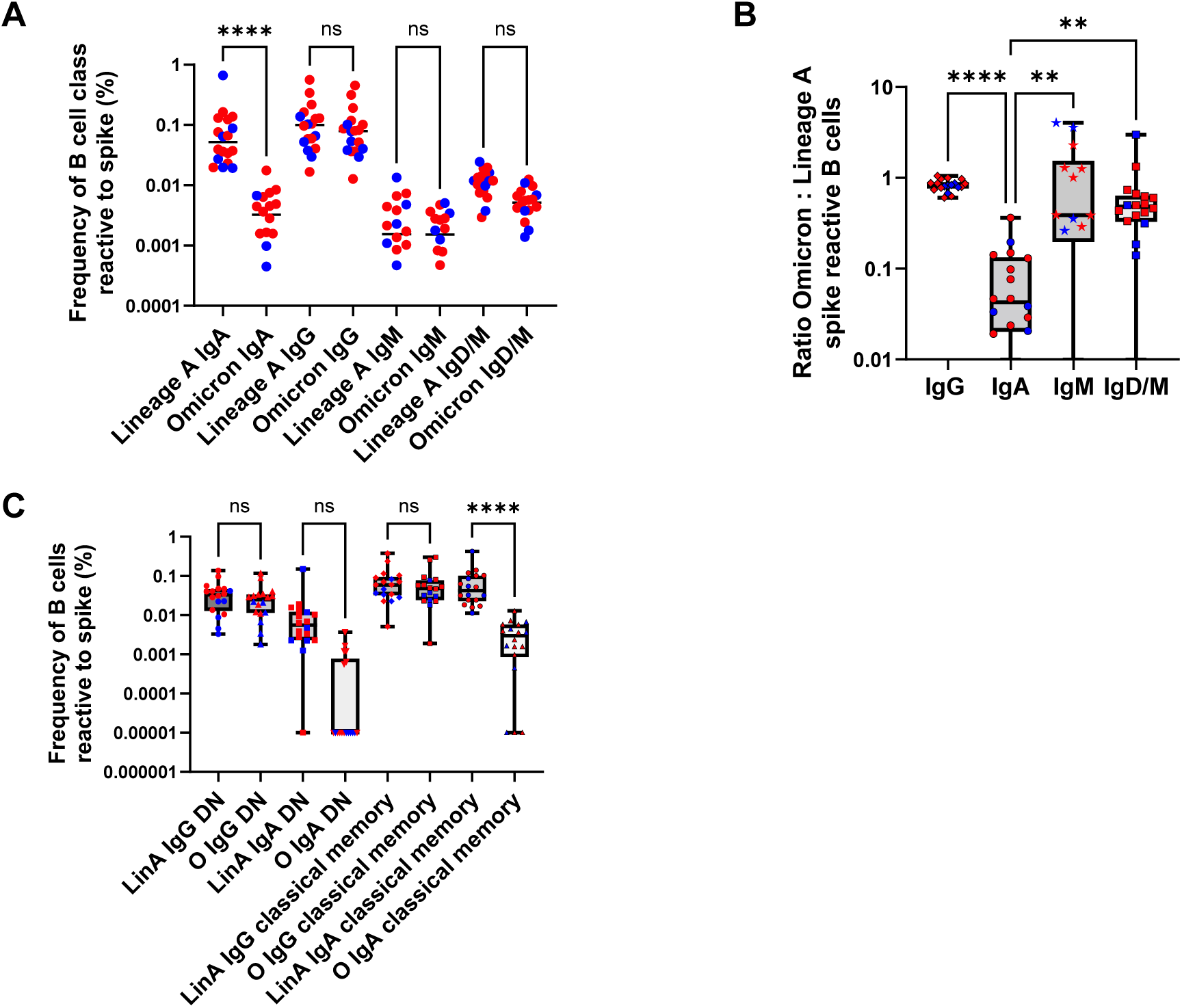
Frequency of B cell subsets reactive to Lineage A and Omicron trimerized spike protein (**A**) Comparison of antibody isotype B cells using flow cytometry showed significantly higher (p<0.0001) frequency of IgA reactive B cells to Lineage A spike compared to Omicron spike. Comparisons of B cells expressing IgG, IgM and IgD/M between Lineage A and Omicron spike did not show any significant differences in the frequency. Blue represents uninfected donors (n=6) whereas red represents previously infected donors (n=12) (**B**) Plot depicting the Omicron:Lineage A ratio of spike reactive B-cells of uninfected (n=6, blue) and previously infected (n=12, red) donors analysed with flow cytometry, showing that the IgG ratio is significantly higher compared to IgA (p<0.0001). The IgM and IgD/M ratios were significantly higher than IgA (p = 0.0029 and p = 0.0059 respectively). (**C**) Comparisons of IgG and IgA double negative (DN) and classical memory B cell subpopulations reactive to Lineage A and Omicron spike of uninfected (n=6, blue) and previously infected (n=12, red) donors. The frequency of Omicron IgA^+^ classical memory B cells was significantly lower than the Lineage A-reactive IgA^+^ classical memory B-cells (p<0.0001). All statistical tests were performed using Prism 9.0 [(**A**) and (**C**) Friedman paired Anova & Dunn’s multiple comparison tests, (**B**) Kruskal-Wallis unpaired Anova & Dunn’s multiple comparison tests].

Having shown in the previous analysis that the frequency of Omicron spike-reactive IgA^+^ B cells is lower than IgG^+^ B-cells, we determined if there were differences in the frequency of non-classical ‘double-negative’ (DN) memory and classical (germinal centre) memory IgG^+^ and IgA^+^ B-cells reactive to Lineage A and Omicron trimeric spike. The double negative (DN) population included DN (CD20^+/-^CD21^+/-^CD24^+/-^CD27^-^CD38^+/-^) and DN2 (CD20^+/-^CD21^-^CD24^-^ CD27^-^CD38^-^) which were both IgD^-^CD27^-^, and the classical memory population encompassed switched activated (CD20^+/-^CD21^-^CD24^-^CD27^+^CD38^-^) and memory switched (CD20^+/-^CD21^+/-^ CD24^+/-^CD27^+^CD38^-^) CD27^+^ B cells. The frequencies of spike-reactive IgG and IgA DN and memory cells were comparable for both Lineage A and Omicron (**Figure 2C**). The median frequency of IgG DN and classical memory cells reactive to Lineage A trimeric spike was 0.0329% (0.0032% – 0.1379%) and 0.0595% (0.0051% - 0.3802%) respectively, whereas for Omicron it was 0.0249% (0.0017% - 0.1163%) and 0.0481% (0.0018% - 0.3017%). In the IgA DN and classical memory cells, the median frequency for Lineage A was 0.0055% (0% – 0.1503%) and 0.0424% (0.0112% - 0.4249%) respectively, whereas for Omicron trimeric spike it was 0% (0% - 0.0037%) and 0.0030% (0% - 0.0129%). Eight uninfected and four infected donors had no detectable IgA^+^ DN B-cells against Omicron spike. Also, one infected and one uninfected donor had no detectable reactive IgA^+^ DN B-cells against either spike. Similarly, three uninfected donors had no detectable reactive IgA^+^ classical memory B-cells against Omicron spike. The frequency of Omicron-reactive IgA^+^ classical memory B-cells was significantly lower (p<0.0001) compared to the respective Lineage A-reactive population (**Figure 2C**). Building on earlier findings that Omicron-specific IgA^+^ B cells are less frequent than IgG^+^ B cells, this analysis further showed that the reduction in IgA^+^ responses extends across both classical memory and non-classical DN B cell subsets. While IgG^+^ classical and DN memory B cells reactive to Omicron and Lineage A were present at similar frequencies, Omicron-reactive IgA^+^ DN and classical memory B cells were significantly reduced, with many donors showing no detectable IgA^+^ B cell reactivity to Omicron at all. This suggests a selective and consistent deficiency in the generation or maintenance of IgA^+^ memory B cell responses against Omicron, potentially impacting mucosal immunity and the ability to neutralize the virus at entry points such as the respiratory tract.

To validate our findings in a more data-driven, unbiased manner, we used weighted unsupervised analysis of spike-specific B cells from the flow cytometry analysis to investigate the dominant B-cell phenotypes reactive to SARS-COV-2 spike and dissect Lineage A and Omicron responses. Ten unique phenotypes were identified by self-organising map analysis (FlowSOM) (**Figure 3A-C**), including four IgA, four IgG/IgA and two IgD/IgM populations. The IgA+ (30%) or IgG+ (49.9%) B cells were all of switched memory (CD19+/CD20+/CD27+) or double negative phenotype (CD19+/CD20+/CD27-). Population 10 as of IgG switched activated memory phenotype, indicated by absence of CD21, whereas all other switched memory populations (CD19+/CD20+/CD21+/CD27+) showed various combinations of CD24, CD27 or CD38 (**Figure 3B**). Each unique B cell population identified by FlowSOM was investigated to understand the frequency cells that were Lineage A or Omicron reactive. There were significantly greater numbers of Lineage A spike reactive B cells in 7/10 of the identified populations (**Figure 3D**). Two of four IgA populations (FlowSOM populations 7 and 8) represented phenotypes that were almost completely devoid of Omicron reactive B Cells (<5%). Levels of Lineage A and Omicron reactive IgG B cell populations (3, 4, 5, 10) were much more similar. Two of the four IgG populations did have lower frequencies of Omicron reactive B cells (population 4, 30.29% Omicron and population 5, 42.46% Omicron), albeit with lower differences than the IgA populations.

**Figure 3.**
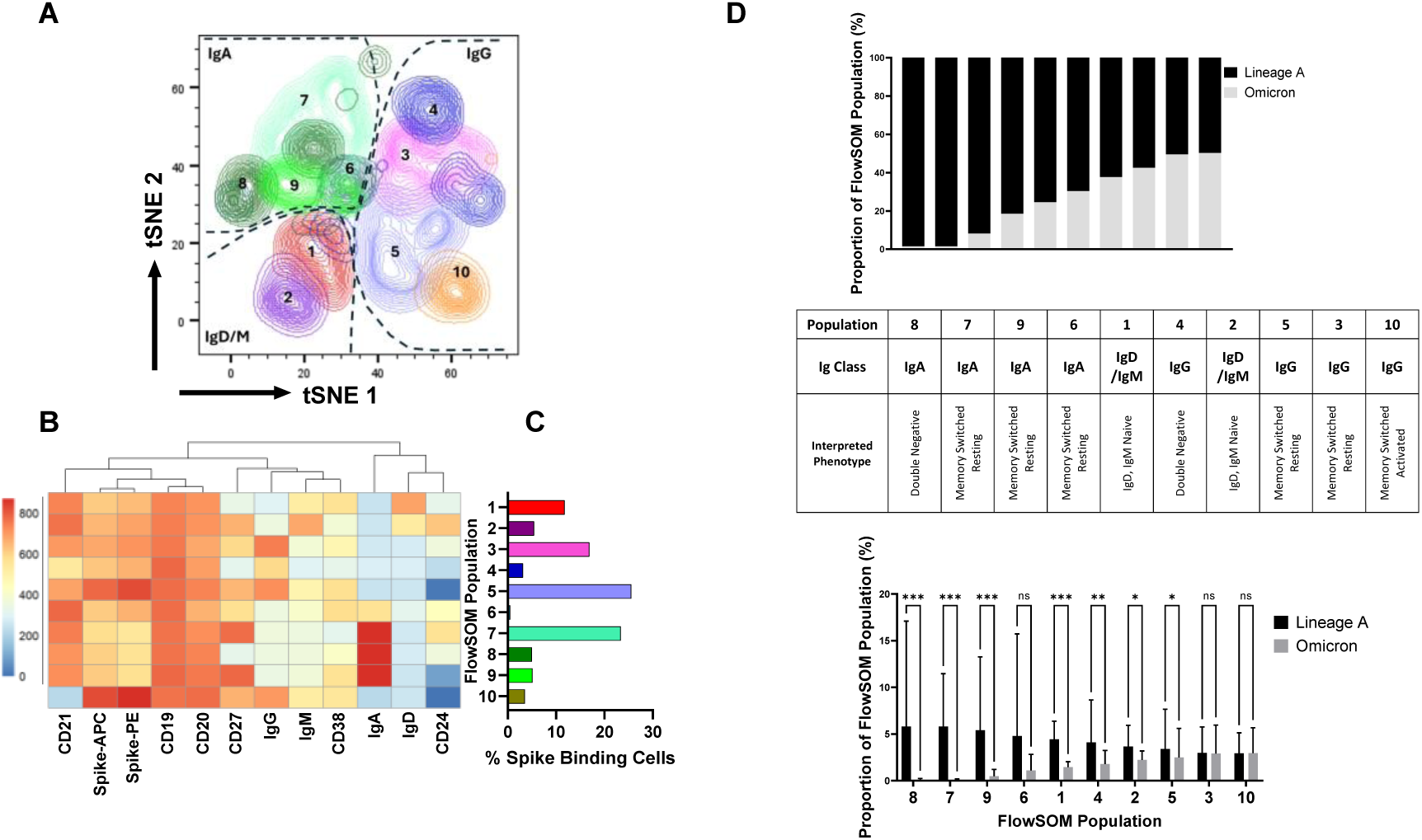
Weighted unsupervised analysis of spike binding B cell phenotypes. Equal numbers of B cells from 17 donors baited with Lineage A or Omicron trimerized spikes were sampled. **(A)** FlowSOM unsupervised analysis identified 10 major populations and are shown on a tSNE representation of the data. **(B)** heatmap of staining intensity among populations. **(C)** Proportion of total spike binding cells by identified population. (D) Proportion of cells binding Lineage A or Omicron spike within a given FlowSOM Populations (upper) and proportion of individual donor Lineage A or Omicron binding cells within each population (lower).

Having analysed the binding and neutralisation potency of donor sera and the frequencies of Ig classes specific to Lineage A and Omicron spike, we next investigated potential correlations amongst these data which could shed light on the relationships between immunity, peripheral B-cells and circulating antibodies. Correlation analysis showed a significant positive correlation between frequency of Lineage A IgG^+^ B cells and Omicron IgG^+^ B cells (p<0.0001) as well as serum binding ED_50_s of Lineage A IgG and Omicron IgG (p<0.0001), suggesting cross-reactivity between the two variants. (**Figure 4A-B**).

**Figure 4.**
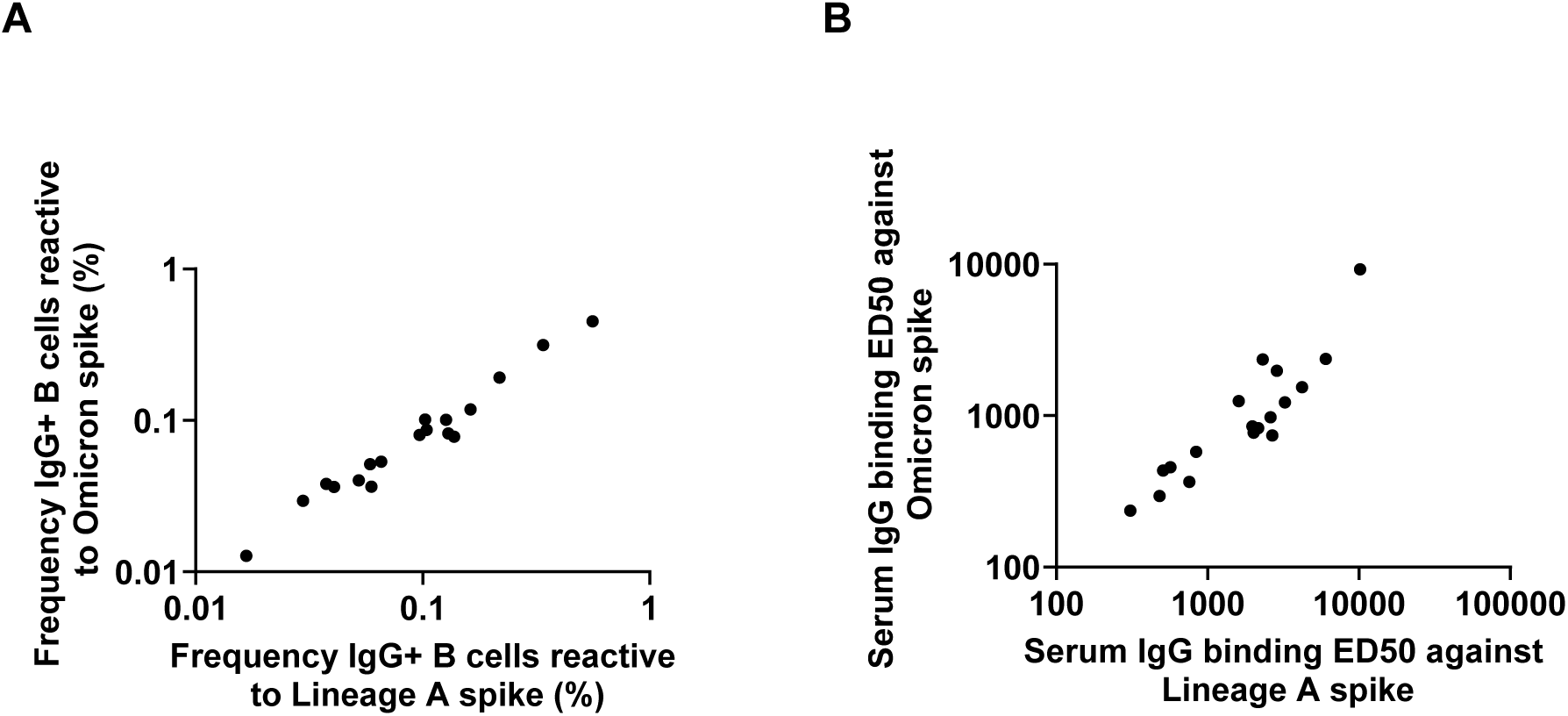
Correlation analysis (non-parametric Spearman) showed significant positive correlations for: (**A**) Frequency of Lineage A IgG^+^ B cells vs frequency of Omicron IgG^+^ B cells (p<0.0001), Data from 2A (**B**) serum binding ED50 Lineage A IgG – ED50 Omicron IgG (p<0.0001), Data from 1B.

### Omicron-reactive B cells in Omicron-naïve donors vaccinated against Lineage A are somatically hypermutated and class-switched

Highly mutated Omicron-binding B cell receptor (BCR) sequences in Omicron-naïve donors would suggest cross-reactivity from previous infections due to memory B cells. To determine whether Omicron-binding B cells arose from mutated B cells, BCR analysis was carried out on a subset of five (n=5) healthcare workers (one uninfected and four previously infected donors) to measure somatic hypermutation (SHM) frequency.

From these samples, 13-154 B cells per donor were sorted by their binding to Lineage A or Omicron trimerized-spikes (**Table 2**). BCR heavy chain sequences were then obtained using PCR amplification and Illumina next-generation sequencing. While both primers were used in this process, far more IgG^+^ BCRs (809) were obtained compared to IgA^+^ (36) with a mean IgA^+^/IgG^+^ ratio of 0.081 across donors and sorts (**Table 3**).

**Table 2.** Cell counts of the number of B cells from the sampled dataset that were sorted and sequenced. BCR sequence counts reflect filtering using different minimum duplicate count thresholds to most closely match the number of cells that were sequenced.

| Donor | Sort | Cell Count | BCR Sequence Count |
| --- | --- | --- | --- |
| **003** | Omicron | 48 | 49 |
| **003** | Lineage A | 13 | 13 |
| **009** | Omicron | 47 | 44 |
| **009** | Lineage A | 47 | 46 |
| **025** | Omicron | 115 | 122 |
| **025** | Lineage A | 93 | 93 |
| **049** | Omicron | 154 | 154 |
| **049** | Lineage A | 73 | 75 |
| **418** | Omicron | 138 | 150 |
| **418** | Lineage A | 94 | 99 |

**Table 3.** Summary of IgA^+^ and IgG^+^ counts and IgA^+^/IgG^+^ ratios per donor and per Lineage A or Omicron sort. , from the sorted B-cell repertoire data.

| Donor | Sort | IgA+ BCR sequences | IgG+ BCR sequences | IgA+/IgG+ Ratio |
| --- | --- | --- | --- | --- |
| **003** | Omicron | 11 | 38 | 0.29 |
| **003** | Lineage A | 0 | 13 | 0 |
| **009** | Omicron | 11 | 33 | 0.33 |
| **009** | Lineage A | 2 | 44 | 0.045 |
| **025** | Omicron | 0 | 122 | 0 |
| **025** | Lineage A | 0 | 93 | 0 |
| **049** | Omicron | 4 | 150 | 0.027 |
| **049** | Lineage A | 8 | 67 | 0.12 |
| **418** | Omicron | 0 | 150 | 0 |
| **418** | Lineage A | 0 | 99 | 0 |

After annotation of V(D)J sequences and identification of clonally related BCRs, we quantified the SHM frequency of each BCR sequence. SHM frequency increases during affinity maturation, the process by which B cells mutate and are selected based on binding affinity to antigens. The presence of mutated BCRs that bind to Omicron without prior exposure to it would indicate the binding of cross-reactive memory B cells, which likely underwent affinity maturation in prior immune responses. We found that IgG BCRs in all donors had an SHM frequency significantly higher than zero (p < 9.9x10^-36^, Z test). Further, in donors 009 and 418, Omicron binding IgG^+^ B-cells were more mutated than those for Lineage A (p < 1.0x10^-166^ and 2.3x10^-40^ for 009 and p < 1.0x10^-166^ and 1.0x10^-166^ for 418, respectively) (**Figure 5**).

**Figure 5:**
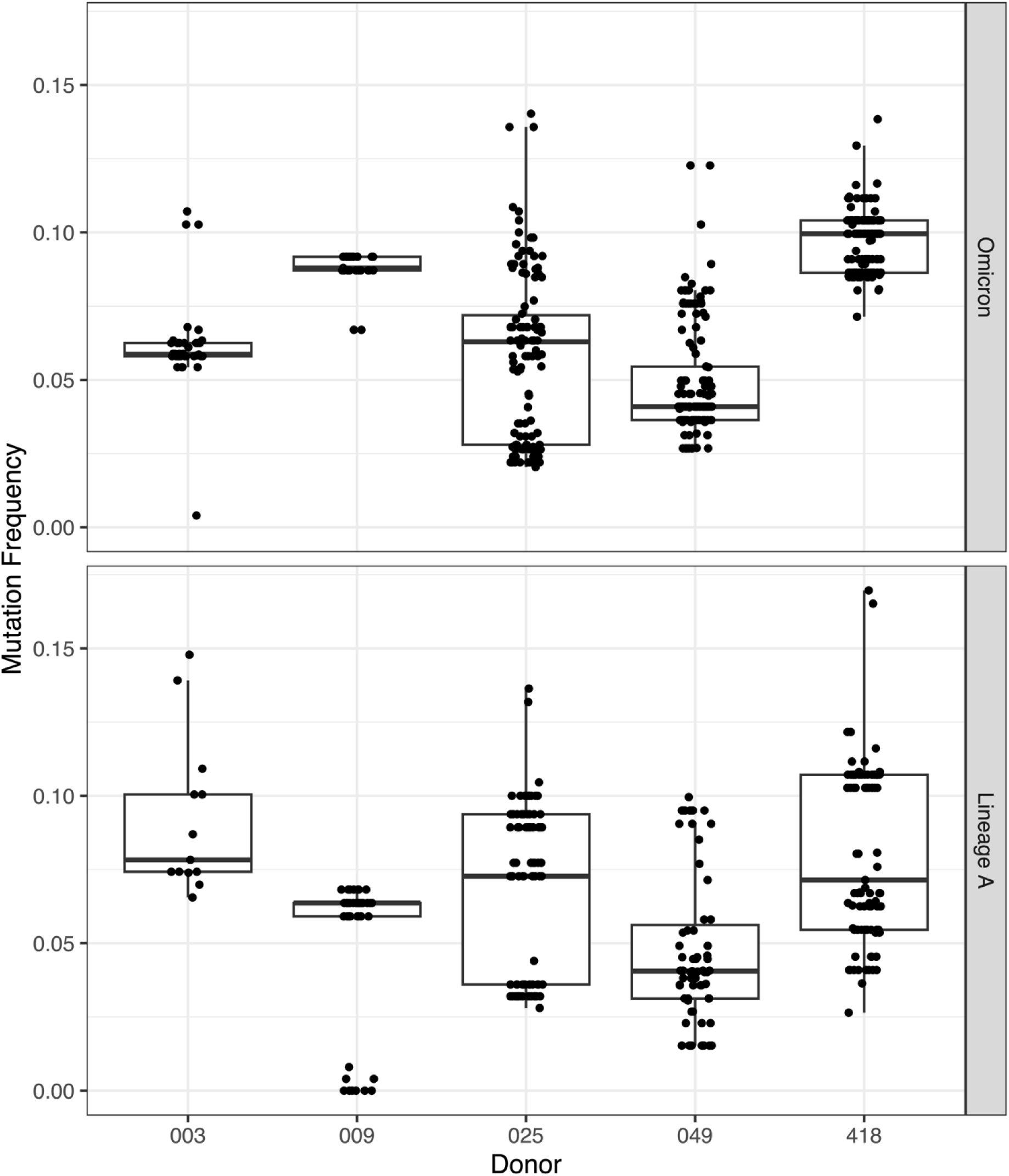
Boxplots showing somatic hypermutation frequency of Omicron and Lineage A binding IgG^+^ B cell sequences for each donor. All values are significant according to a single-sample Z-test against a null hypothesis of 0 mutations. In donors 009 and 418, Omicron binding IgG^+^ B-cells were even more mutated than those for Lineage A.

Multiple studies have identified highly similar SARS-CoV-2 binding antibodies across donors, indicative of common antibody responses [18]. To determine whether the IgG BCRs obtained were similar to previously published Lineage A and Omicron binding antibodies, we compared the BCR data to the publicly available CoV-AbDab database of previously published SARS-CoV-2 binding antibodies [19]. Within our dataset, we identified 123 sequences from 7 clones that had the same V gene, J gene, CDR3 length, and >80% CDR3 amino acid similarity to sequences in the CoV-AbDab. Three of these clones matched to sequences that were both Omicron and Lineage A neutralizing, three used the IGHV3-30 V gene, two were exclusively Lineage A neutralizing, and one had both IgA^+^ and IgG^+^ isotypes (**Table 4**). The largest of these clones (114) that neutralizes both Omicron and Lineage A aligned well with four previously published antibody sequences (BD56-556, BD56-657, PDI-222, BD56-1872) in the CoV-AbDab database (**Figure 6**). This indicates that some of the Omicron-binding IgG^+^ B cells that were found were highly similar to previously published Omicron and Lineage A cross-reactive antibodies. Overall, these results demonstrate that Omicron-binding IgG^+^ B cells in donors naïve to Omicron but vaccinated against Lineage A were mutated and class-switched. This implies these Omicron-binding B cells were antigen-experienced, and likely cross-reactive memory B cells from Lineage A vaccination and/or infection with a variant other than Omicron.

**Figure 6:**
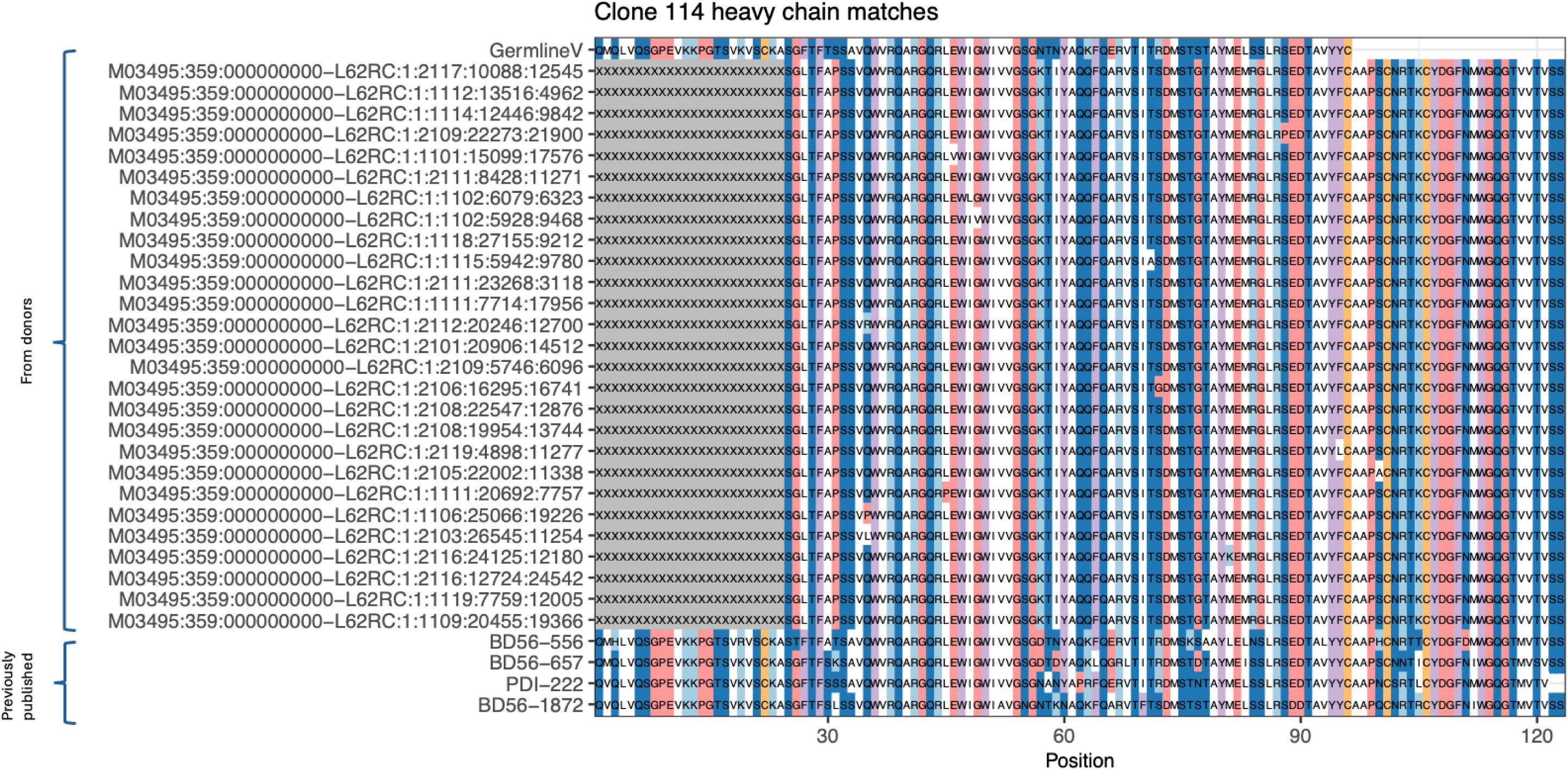
Amino acid multiple sequence alignment of convergent sequences in clone 114, the largest Omicron neutralizing clone, and previously published sequences in the CoV-AbDab database (antibodies BD56-556, BD56-657, PDI-222, BD56-1872). Clone 114 used V genes IGHV1-58*01 and IGHV1-58*03. IgG^+^ B cell sequences bind to the receptor binding region of the spike protein of both variants (Lineage A and Omicron) of SARS-CoV-2 and neutralize them. The clone had an average mutation frequency of 0.090. See also Table 4.

**Table 4.** Summary of seven clones with convergent sequences with the public CoV-AbDab database of SARS-CoV-2 antibodies.

| Clone | Donor | Sort | V gene | Matches from CoV-AbDab | Epitopes | Strains neutralized | Isotype | Mean SHM frequency |
| --- | --- | --- | --- | --- | --- | --- | --- | --- |
| **114** | 009 | O | IGHV1-58*01, IGHV1-58*03 | BD56-556, BD56-657, PDI-222, BD56-1872 | S, RBD | SARS-CoV2_WT, SARS-CoV2_Omicron-BA1, SARS-CoV2_Omicron-BA1.1, SARS-CoV2_Omicron-BA3, SARS-CoV2_Omicron-BA2, SARS-CoV2_Omicron-BA2.12.1, SARS-CoV2_Omicron-BA2.13, SARS-CoV2_Omicron-BA4/5, SARS-CoV2_Omicron-BA2.75, SARS-CoV2_Omicron-BA5, SARS-CoV2_Omicron-BA4/5 (weak), SARS-CoV2_Omicron-BA5 (weak) | IgG | 0.09 |
| **17** | 025 | O | IGHV2-5*01 | BD55-1472, BD56-622 | S, RBD | SARS-CoV2_WT (weak), SARS-CoV1, SARS-CoV2_Omicron-BA2.12.1 (weak), SARS-CoV2_Omicron-BA1, SARS-CoV2_Omicron-BA2, SARS-CoV2_Omicron-BA2.75, SARS-CoV2_Omicron-BA5, SARS-CoV1 (weak), SARS-CoV2_Omicron-BA1 (weak), SARS-CoV2_Omicron-BA1.1 (weak), SARS-CoV2_Omicron-BA3, SARS-CoV2_Omicron-BA2 (weak), SARS-CoV2_Omicron-BA2.13 (weak), SARS-CoV2_Omicron-BA4/5 (weak), SARS-CoV2_Omicron-BA2.75 (weak), SARS-CoV2_Omicron-BA5 (weak), SARS-CoV2_Omicron-XBB (weak) | IgG | 0.039 |
| **128** | 418 | LinA | IGHV1-58*02 | BD57-022, BD56-447, ZCB11, BD57-0385 | S, RBD | SARS-CoV2_WT, SARS-CoV2_Omicron-BA1, SARS-CoV2_Omicron-BA1.1, SARS-CoV2_Omicron-BA3, SARS-CoV2_Omicron-BA2, SARS-CoV2_Omicron-BA2.12.1, SARS-CoV2_Omicron-BA2.13, SARS-CoV2_Omicron-BA4/5 (weak), SARS-CoV2_Omicron-BA2.75, SARS-CoV2_Omicron-BA5 (weak), SARS-CoV2_Alpha, SARS-CoV2_Beta, SARS-CoV2_Gamma, SARS-CoV2_Delta, SARS-CoV2_Omicron-BA4.6 (weak), SARS-CoV2_Omicron-BA4.7 (weak), SARS-CoV2_Omicron-BA5.9 (weak), SARS-CoV2_Omicron-BL1, SARS-CoV2_Omicron-BN1, SARS-CoV2_Omicron-BA2.75.1, SARS- | IgG | 0.041 |
|  |  |  |  |  |  | CoV2_Omicron-BA2.75.4, SARS-CoV2_Omicron-BA2.75.5, SARS-CoV2_Omicron-BA2.75.6 |  |  |
| **36** | 049 | O | IGHV1-18*01,<br>IGHV1-18*04 | 368.13.C.0123 | S, Unk |  | IgG, IgA | 0.04 |
| **43** | 049 | LinA | IGHV3-30*18,<br>IGHV3-30-5*01 | ADI-70080, Turner-22-1B04 | S, RBD |  | IgG | 0.054 |
| **45** | 049 | O | IGHV3-30*01,<br>IGHV3-30*10 | S2B5,<br>S2B7,<br>S2H12,<br>S2110#,<br>368.07.C.0103,<br>368.07.C.0219,<br>368.10.D.0032,<br>368.10.D.0114,<br>368.10.D.0143,<br>P054_045,<br>PDI-196,<br>R121-3C12, R259-1A4, R259-3D1,<br>R410-3A10, R410-3G12, R616-1A9,<br>WCSL-44, WCSL-70 | S, S2, Unk,<br>non-S1, non-RBD | SARS-CoV2_WT, SARS-CoV2_WT (weak) | IgG | 0.032 |
| **47** | 049 | LinA | IGHV3-30*18,<br>IGHV3-30-5*01 | S2B5,<br>S2H12,<br>368.10.D.0171,<br>R259-3F2,<br>WCSL-44 | S, S2, Unk,<br>non-RBD | SARS-CoV2_WT | IgG | 0.058 |

## Discussion

Our findings demonstrate that Omicron-reactive IgG^+^ memory B cell and antibody responses are preserved in Omicron-naïve individuals who were previously vaccinated with Lineage A and/or infected with a pre-Omicron SARS-CoV-2 variant. In contrast, IgA^+^ B cell responses are markedly diminished, suggesting a potential impairment in mucosal immunity to Omicron, even in those with prior SARS-CoV-2 infection. This contrasts with a recently published study showing elevated levels of Lineage A spike-specific mucosal IgA in previously infected and vaccinated individuals compared to those without prior infection [20]. Early SARS-CoV-2 specific humoral responses are dominated by IgA antibodies, which contribute more significantly to virus neutralisation than IgG [21, 22]. Although IgA antibodies are primarily found at mucosal surfaces, previous studies have demonstrated strong clonal relationships between peripheral and mucosal IgA responses [23]. Moreover, serum and mucosal IgG levels correlate strongly, likely due to IgG spillover from the periphery to mucosal sites [20, 24]. Trimeric spike-specific IgA titres decline within a month after symptom onset; however, neutralising IgA can persist in saliva for longer periods [21, 25]. A recent study has shown that nasal IgA responses wane several months after infection and are minimally boosted by subsequent vaccination [26]. This aligns with our findings of Omicron-reactive IgA^+^ memory B cells compared to Lineage A, potentially explaining the limited protection against reinfection and the modest impact of vaccination on transmission.

Our data also shows that vaccination and/or infection generate both serum IgG and IgG^+^ memory B cells that can recognise Omicron spike, but this cross-reactivity is severely reduced in the peripheral IgA^+^ memory B cells. The IgG reactivity against Lineage A and Omicron spikes and the proportion of IgG^+^ B cells reactive to both spikes were equivalent. Similarly, the levels of IgG^+^ DN and classical memory populations were equivalent against both variants. Several studies have shown that vaccination with ancestral Lineage A SARS-CoV-2 induce durable immune memory to Lineage A and variants of concern including Omicron [27, 28], although mucosal IgA is reportedly blunted [29]. Also, a study in macaques showed that boosting with Omicron spike elicits similar B cell protection as boosting with Lineage A [30]. As mentioned above, our study shows equivalent levels of IgG^+^ DN memory B cells for both variants and a trend towards a lower frequency of IgA^+^ DN memory B cells against Omicron. The atypical extrafollicular memory B cell populations in our study included DN (CD20^+/-^CD21^+/-^CD24^+/-^CD27^-^CD38^+/-^) and DN2 (CD20^+/-^CD21^-^CD24^-^ CD27^-^CD38^-^). However, DN1 (CD27^-^CD38^+^CD24^+^CD21^+^) were only observed in three donors. Extrafollicular activation is strongly correlated with large antibody-secreting cell expansion and early production of high concentrations of SARS-CoV-2-specific neutralising antibodies [31].

Vaccination provides immunity against SARS-CoV-2 by generating bone-marrow-resident antibody secreting cells, giving rise to circulating neutralising antibodies, as well as memory B cells, which can rapidly respond upon re-exposure to antigen. Previous studies have shown that immune protection, especially against infection is reduced for Omicron compared to other VOCs. [3–6, 32]. A recent longitudinal study showed that individuals who recovered from Lineage A natural infection retained immunity for two years post-infection, however, neutralising antibody responses were diminished against newly emerged variants [33]. Our study shows that although both peripheral spike-specific IgG levels and IgG^+^ memory B cell frequencies are similar for Lineage A and Omicron in vaccinated donors with or without previous infection, the serum neutralisation potency against Omicron pseudotypes is decreased compared to Lineage A. Neutralisation is one mechanism by which antibodies provide protection, other protective mechanisms include Fc-dependent effector functions (reviewed in [34]) such as antibody-dependent cellular cytotoxicity (ADCC) [35], antibody-dependent complement deposition (ADCD), antibody-dependent cellular phagocytosis (ADCP), and opsonisation [36], and it will be important to define whether the Omicron-specific non-neutralising antibodies can exert these effects.

Our BCR analysis implied the presence of memory B cell clones cross-reactive between Lineage A and Omicron. We did not directly find clonal overlap between strains as evidence for cross-reactivity between them. However, this may be due to the small sample of cells that were sequenced and sorted. The pre-Omicron presence of highly mutated Omicron-binding BCRs, as well as public antibody matches (S2B5, S2H12, and WCSL-44) between two Omicron and Lineage A sorted clones (45 and 47 in **Table 4**) from the same donor (49) supports the conclusion of cross-reactive memory B cells. Additionally, a Lineage A sorted clone (128 in **Table 4**) had antibodies matching previously published Omicron-neutralising ones.

In summary, while vaccination or prior infection with Lineage A SARS-CoV-2 induces cross-reactive and somatically mutated IgG^+^ memory B cells capable of recognizing Omicron, the markedly diminished frequency of Omicron-specific IgA^+^ memory B cells suggests a potential vulnerability to Omicron infection at mucosal entry points. However, the presence of Omicron-reactive IgG^+^ memory B cells may still contribute to systemic immune protection and reduce the risk of severe disease, even in the absence of robust mucosal IgA responses. These insights were made possible by studying a unique cohort of healthcare workers who had been previously infected and/or vaccinated against Lineage A but had not been exposed to the antigenically distinct Omicron variant prior to sample collection, allowing clear delineation of cross-reactive immune memory.

## Materials and methods

### Study design

The PANTHER study is a longitudinal cohort study launched in April 2020 to follow up seropositivity to SARS-CoV-2 among healthcare workers in Nottingham, United Kingdom. The cohort is described in detail in [37] and [17].

Participants who had taken part in the previous serological surveillance study were invited to take part in the post-boost vaccination study. Twelve participants with a history of infection, as determined by the Roche N ELISA and six individuals without infection were included in the present study (**Table 1**). Sixteen (n=16) of them have had 3 doses of BNT162b2 and two (n=2) have had 2 doses of ChAdOx1-S & 1 dose of BNT162b2. The first vaccine dose was administered in December 2020/January 2021, the second in February/March 2021 and the booster in September/October 2021. The healthcare workers took weekly lateral flow assay tests and none tested positive in the period between the introduction of Omicron into the UK and the time of sampling used in this study, *i.e.,* the volunteers were Omicron naïve. Plasma and PBMCs were collected in January 2022, ∼106 days on average (ranging from 70 to 126 days) since the donors received their booster dose, to assess their responses against Lineage A and Omicron variants. Samples were collected and stored under a Human Tissue Authority (HTA) license in the Nottingham Tissue Bank (license number 11035). The study protocol was approved by Northwest–Greater Manchester South Research Ethics Committee (reference 20/NW/0395). All assays were performed in duplicate (serology) or triplicate (neutralisation).

### Donor sampling

Thirty (30) millilitres of blood were drawn from each donor. Peripheral blood mononuclear cells (PBMC) and plasma were collected after processing the blood samples with Histopaque-1077 (Sigma-Aldrich) and SepMate PBMC isolation tubes (Stemcell Technologies). The plasma samples were heat inactivated at 56°C for 30 min. The samples were stored according to the HTA guidelines.

### Spike-specific enzyme immunoassay

Lineage A SARS-CoV-2 trimeric spike glycoprotein manufactured in CHO cells and Omicron SARS-CoV-2 trimeric spike glycoprotein manufactured in HEK293 cells were obtained from The Native Antigen Company (REC31871 and REC32008 respectively). Recombinant proteins were immobilised onto microtiter plate wells and used to capture spike-specific antibodies present in indicated dilutions of human serum samples. Captured IgG and IgA were detected using a human IgG–specific and IgA-specific horseradish peroxidase–conjugated antibody respectively, followed by the addition of ultra-3,3′,5,5′-tetramethylbenzidine substrate (Thermo Fisher Scientific). Absorbance at 450 nm was read using a FLUOstar Omega plate reader (BMG Labtech). Data were presented as the ratio of the absorbance (OD_450_) for the sample versus the background absorbance. Binding curves were generated using heat-inactivated human plasma starting at 1/20 dilution down to 1/81,920 (**Supplementary** Figure 1). The ED_50_ value for each sample was calculated using a non-linear analysis model on GraphPad Prism version 9.3.1 (Graphpad Software, San Diego, USA).

### Pseudotype neutralisation assay

Pseudotype neutralisation assays were performed as previously described [17] and [38]. Neutralisation curves were generated using heat-inactivated human plasma starting at 1/100 dilution down to 1/62,500 (**Supplementary** Figure 2). The ID_50_ value for each sample was calculated using a non-linear analysis model on GraphPad Prism version 9.3.1 (Graphpad Software, San Diego, USA).

### Flow cytometry analysis

Ten million (10 x 10^6^) PBMCs were defrosted for each donor and incubated O/N in Gibco™ RPMI 1640 Medium + 10% fetal bovine serum (FBS). The cells were washed with 3% bovine serum albumin (BSA) in PBS and each sample was divided into two; one for Lineage A and one for Omicron spike. The cells were blocked with TruStain FcX ‘FC Block’ for 15min at 4°C and then stained with the antibody mix (**Supplementary Table1**) for 30min at 4°C. After washing again, the cells were stained with biotinylated Lineage A or Omicron trimeric spike (REC31871 and REC32008 respectively ,The Native Antigen Company) for 30min at 4°C followed by washing and staining with R-PE and APC conjugated streptavidin (Thermo Fisher Scientific) for 30min at 4°C. After the final wash, the cells were stained with Annexin efluor 450 and propidium iodide (PI) (Thermo Fisher Scientific) for 20min at 4°C. The samples were analysed on a ID7000 Spectral Flow Cytometer (SONY).

### Statistical analysis

All statistical analyses were performed using GraphPad Prism version 9.3.1 unless otherwise noted. Tests used are noted in the appropriate figure legends.

### Weighted unsupervised analysis

Single, viable (Annexin V-/PI-) lymphocytes were identified and B cells further gated based on CD19 and CD20 expression (FlowJo Version 10). Gating shown in **Supplementary** Figure 3. Equal numbers of these viable single B cells from paired samples baited with Lineage A or Omicron trimerized spike were down-sampled (DownSample V3 plugin, 59K per sample) and within these spike-R-PE+/spike-APC+ (double positive) cells were gated and entered into an unsupervised self-organising map analysis (FlowSOM, using R plugin for FlowJo)[39]. A tSNE plot was generated for data visualisation using opt-SNE [40] (settings: iterations 1000, perplexity 30, learning rate 279, Barnes-Hutt gradient algorithm and exact (vantage point tree) KNN algorithm).

### B cell receptor sequence repertoire processing and analysis

From the donor samples, 13-154 B cells from 5 donors were sorted by binding to Lineage A or Omicron trimerized-spikes, amplified using PCR targeting the V-genes and the constant regions for IgG and IgA, and sequenced using Illumina 2x250bp paired-end next-generation sequencing to produce a repertoire of B cell receptor (BCR) sequences.

The BCR sequence data were pre-processed using pRESTO v0.7.2 [41] and analysed using Python v3.12.2 and R v4.3.3 packages from the Immcantation (www.immcantation.org) framework. Paired-end reads were assembled into sequences using mate-pair alignment. Initial quality control was performed by removing all reads with a Phred quality score < 20. V-segment primers were aligned with the reads to ensure their orientation in the direction of the V(D)J reading frame. The C-region primer was aligned to each read to identify antibody isotype. For both alignment steps, primers were scanned across the full length of each assembled sequence with a maximum allowable error rate of 0.1. Since primer regions can have high error rates, the V-segment and C-region primer regions were masked for each assembled sequence. Duplicate sequences, sharing the same isotype, were collapsed to a single unique sequence while allowing for up to 20 interior N-valued positions and within groups defined by V-segment primer annotations. The sequences were filtered to those with at least 2 duplicates and a maximum of 20 ambiguous characters. V and J gene annotations for each sequence were obtained by alignment to the IMGT GENE-DB human reference database [42] using IgBlast v1.22.0 [43] within the Immcantation Docker container v4.5.0.

For each strain from each donor, the repertoire was filtered by choosing a minimum duplicate count threshold to most closely match the number of B cells that were sorted. Sequences were assigned to clones using single-linkage hierarchical clustering among sequences with the same V gene and J gene annotation, same junction length, and at most 10% nucleotide difference in the junction using the SCOPer v1.3.0 [44] package. The unmutated germline sequence for each clone was reconstructed using Dowser v2.2.1 [45] using the same IMGT database used earlier. Somatic hypermutation frequency was calculated for each sequence based on the length-normalized Hamming distance along the V gene (IMGT positions 1-312) using SHazaM v1.2.0 [46]. To determine whether the B cells in a sample had mutation frequencies significantly greater than zero, a single sample Z-test was carried out using BSDA v 1.2.2 [47] against a null hypothesis of 0 mutations. To identify BCRs with convergent antibody sequences, all sequences were compared to the publicly available CoV-AbDab database (obtained 06/08/2024) containing 12,916 previously published SARS-CoV-2 binding antibodies [19]. BCRs were considered convergent if they had the same V and J gene annotations, CDR3 length, and at most 20% CDR3 amino acid difference with a heavy chain in the CoV-AbDab. Results visualized using ggpubr3 v0.6.0 [48]. Scripts for replicating B cell receptor sequence repertoire processing and analysis are available at https://github.com/hoehnlab/publication_scripts.

---

**Supplementary Figure 1.**
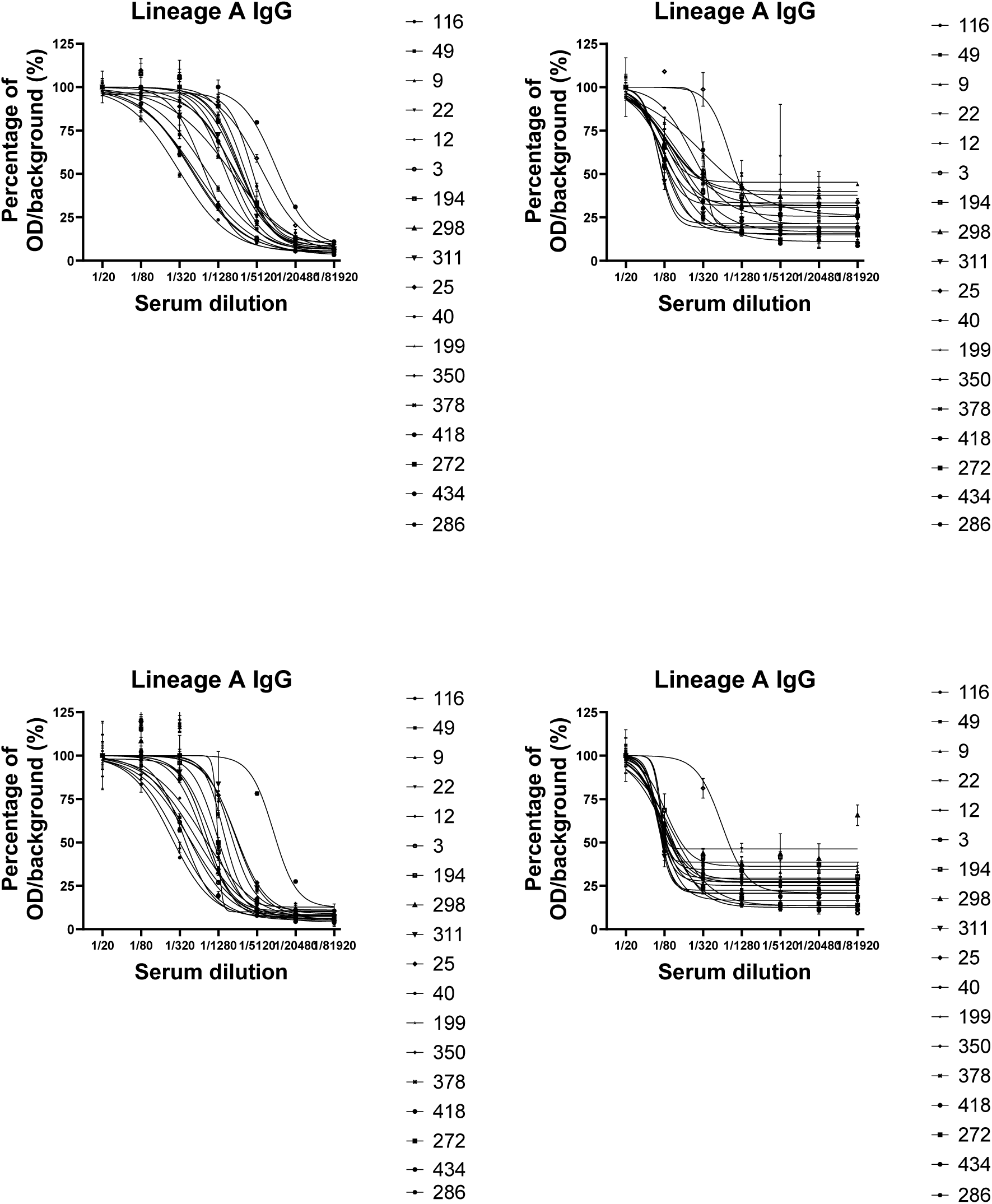
Binding curves of heat-inactivated human plasma IgG and IgA to SARS-CoV-2 Lineage A and Omicron trimeric spike glycoprotein. The dilution series of the human plasma started from 1 in 20 down to 1 in 81,920.

**Supplementary Figure 2.**
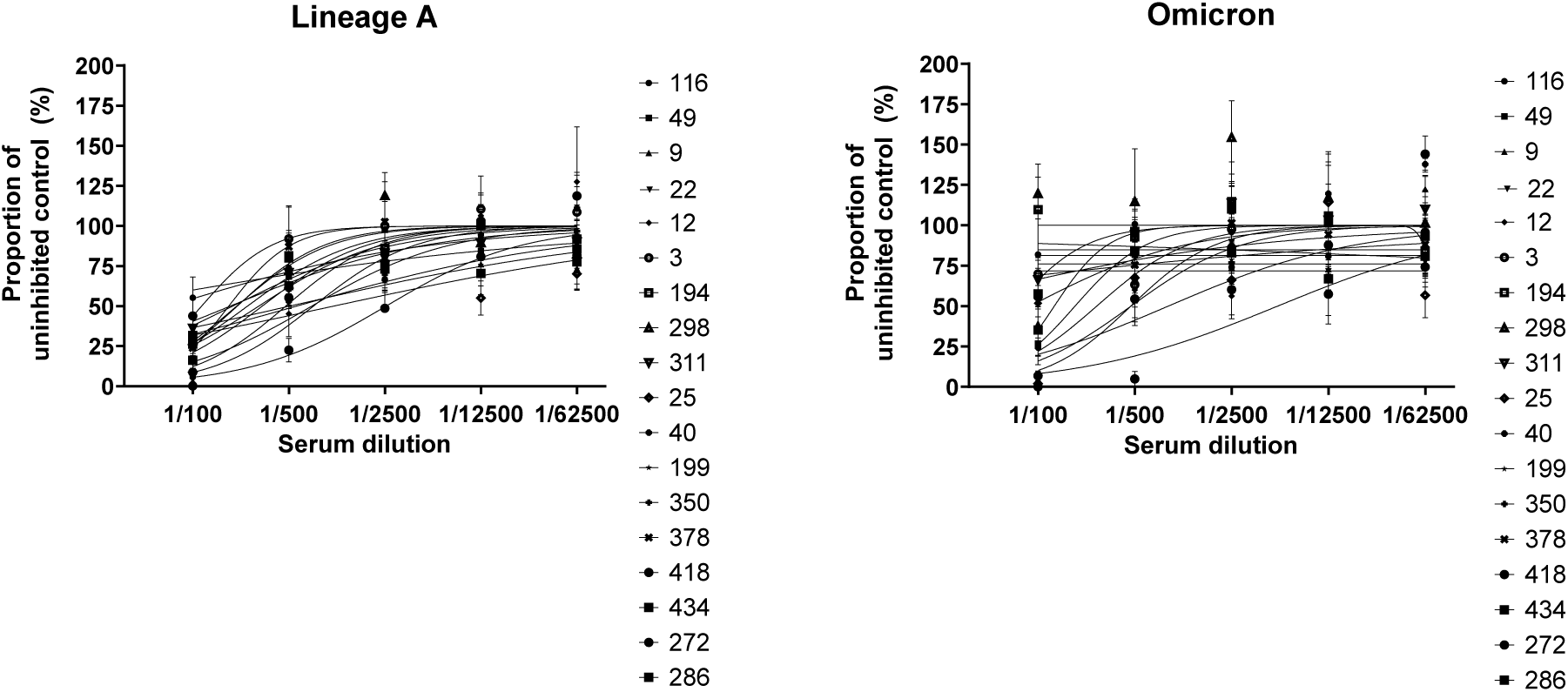
Neutralisation curves of heat-inactivated human plasma against SARS-CoV-2 Lineage A and Omicron pseudotypes. The dilution series of the human plasma started from 1 in 100 down to 1 in 62,500.

**Supplementary Figure 3.**
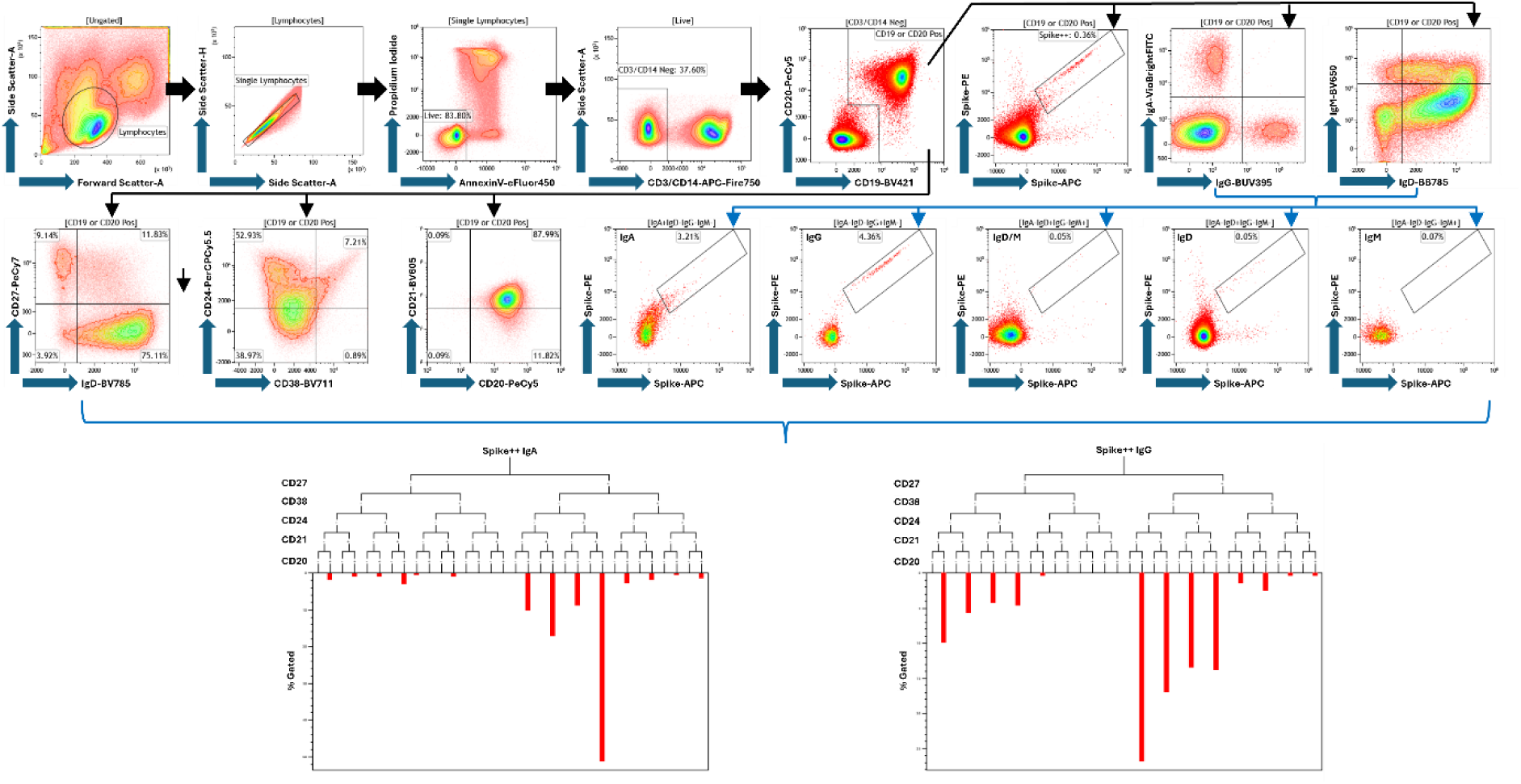
Example Gating of Flow Cytometric Analysis. Viable single lymphocytes were gated and CD3+ or CD14+ cell removed before a total B cell gate of CD19 or CD20 was identified. Total B cells were then examined for expression of IgA, IgD, IgG and IgM as well as phenotype markers CD21, CD27 and CD38. Spike++ events were identified in total B -cells as well as gated immunoglobulin class populations. Immunoglobulin class and phenotyping gates were combined to explore phenotypic combinations in Spike ++ events using Tree-plots (IgA and IgG Shown). Example data shown is donor 25 bated with lineage A spike.

**Supplementary table 1.** List of the markers and fluorophores used in the flow cytometry panel to characterise B cell populations within human PBMC.

|  | Fluorophore | Ex (nm)<br>Max | Em (nm)<br>Max | Clone | Product Number | Supplier |
| --- | --- | --- | --- | --- | --- | --- |
| Spike | PE | 565 | 575 |  |  |  |
| CD20 | PE/Cyanine5 | 565 | 670 |  |  |  |
| Spike | APC | 650 | 660 |  |  |  |
| CD19 | Brilliant Violet 421™ | 405 | 421 |  |  |  |
| CD38 | Brilliant Violet 711™ | 405 | 711 | HIT2 | 303528 | Biolegend |
| CD27 | PE/Cyanine7 | 565 | 774 |  | 302838 | Biolegend |
| CD21 | Brilliant Violet 605™ | 405 | 603 | Clone B-ly4 | 740395 | BD |
| IgM | Brilliant Violet 650™ | 405 | 645 | MHM-88 | 314526 | Biolegend |
| IgD | Brilliant Violet 785™ | 405 | 785 | Clone IA6-2 | 348242 | Biolegend |
| CD3/CD14 | APC/Fire™ 750 | 650 | 774 |  | 300470 AND 367120 | Biolegend |
| CD24 | PerCP/Cyanine5.5 | 482 | 690 | ML5 | 311116 | Biolegend |
| IgG | BUV 395 |  |  | G18-145 | 564229 | BD |
| IgA | Vio Bright FITC | 493 | 525 | IS11-8E10 | 130-113-480 | Miltenyi |
| Annexin V | eFluor450 |  |  |  |  |  |
| L/D | PI |  |  |  |  |  |

